# MMonitor: Software for Real-Time Monitoring of Microbial Communities Using Long Reads

**DOI:** 10.1101/2025.09.08.674848

**Authors:** Timo N. Lucas, Ulrike Biehain, Anupam Gautam, Kurt Gemeinhardt, Tobias Lass, Simon Konzalla, Ruth E. Ley, Largus T. Angenent, Daniel H. Huson

## Abstract

Real-time monitoring of microbial communities offers valuable insights into microbial dynamics across diverse environments. However, many existing metagenome analysis tools require advanced computational expertise and are not designed for monitoring. We present MMonitor, an open source software platform for the real-time analysis and visualization of metagenomic Oxford Nanopore Technologies (ONT) sequencing data. MMonitor includes two components: a desktop application for running bioinformatics pipelines through a graphical user interface (GUI) or command-line interface (CLI), and a web-based dashboard for interactive result inspection. The dashboard provides taxonomic composition over time, quality scores, diversity indices, and taxonomy–metadata correlations. Integrated pipelines enable automated de-novo assembly and reconstruction of metagenome-assembled genomes (MAGs). To validate MMonitor, we tracked human gut microbial populations in three bioreactors using 16S rRNA gene sequencing, and applied it to whole-genome sequencing (WGS) data to generate high-quality annotated MAGs and reveal functional insights. We also compare MMonitor to other software for real-time metagenomic analysis, highlighting the strengths and limitations of each tool for this use case. By performing automated time-series analyzes, sample management, and updating of reference databases, MMonitor addresses current limitations and supports dynamic microbiome research in various fields.

**IMPORTANCE:** Metagenome monitoring is essential for understanding and managing microbial communities in dynamic environments. Rapid and accurate analysis of these communities allows for timely interventions in biologically active environments, for example, in industrial biotechnology, environmental surveillance, and medical diagnostics. Metagenome monitoring tools should have real-time capabilities, intuitive interfaces, convenient setup and distribution, and suitable visual representations for effective monitoring. MMonitor addresses these challenges by offering a platform that combines real-time analysis of nanopore sequencing data with customizable pipelines and advanced visualization tools. It is designed to be accessible to researchers of all levels of computational skill. It supports efficient taxonomic and functional analyses based on popular metagenome software and automates taxonomic profiling, genome assembly, binning, and annotation. The dashboard helps to interpret data and present results. MMonitor is designed to track metagenomes and we hope that it can impact microbiome research and applications in health, biotechnology, agricultural, and environmental sciences.

## INTRODUCTION

### Monitoring metagenomes through long-read sequencing

The study of microbial communities through metagenomics has advanced our understanding of microbial diversity and function in various environments [1]. Microbiome sequencing and bioinformatic analysis are used to monitor bioactive systems, such as anaerobic digesters, to optimize their performance, or human microbiome systems, to study host-microbiome interactions and their effects on human health. [2–4]. Metagenomic monitoring also connects microbial dynamics with biogeochemical processes in larger-scale ecosystems, such as the ocean or the soil [5, 6].

However, while optical, fluorescence-based, and qPCR assays can sensitively quantify specific taxa, they are limited to predefined targets, making it difficult to capture the overall dynamics of complex microbial communities. Long-read sequencing technologies, such as ONT platforms, have emerged as powerful tools in this field. Compared to short-read sequencing, long-read technologies offer higher taxonomic resolution, simplifying the assembly of complete reference genomes from metagenomes or pure cultures, without the need for complementary short-read data [7, 8]. High-quality genomes are essential for detecting structural variations in microbial genomes [9] and improving annotations to assess the genetic potential of microbial communities [10].

The analysis of long-read data poses unique computational challenges. Algorithms designed for short read data often struggle with the higher error rates and sequence lengths. Efficient algorithms are essential for real-time monitoring, particularly in time-sensitive settings such as pathogen detection, where rapid analysis can aid in interventions and decision making. New algorithms for taxonomic and functional analysis of long reads have emerged to address this challenge. Tools such as MetaMaps [11] and Emu [12] which were specifically designed for long-read data, while others, such as Centrifuge and Kraken2, originally built for short-read analysis, have also proven useful for long-read metagenomics [13].

Nanopore sequencing has been successfully applied in various real-time monitoring applications, such as tracking microbial resistance genes [14], identifying disease outbreaks [15], providing reference-agnostic surveillance of pathogens in insects [16], or freshwater environments [17]. The real-time analysis capability of nanopore sequencing is useful for large-scale industrial applications, where undetected contaminations could lead to costly production losses. The MinION device from ONT, with its portability and short sample-to-sequence time, has been deployed beyond traditional laboratory settings, including the International Space Station [18] and ocean research vessels [19].

Although several tools have implemented real-time monitoring, such as minoTour [20], NanopoReaTA [21], MARTi [22], and Nanometa Live [23], these tools face challenges in scalability, ease of use, and flexibility. Many focus heavily on taxonomic profiling without a real-time component, and others lack features, time-series visualization, or effective sample management. Metagenomic analysis pipelines, such as HUMAnN [24] or QIIME [25], can generate comprehensive reports, but these tools can require command-line expertise, limiting their accessibility. Online platforms, such as MG-RAST [26], BusyBee [27] and BugSeq [28], provide user-friendly interfaces. However, they do not offer an analysis of continuous sequencing data streams. Furthermore, without consistent updates, the monitoring software suffers from a reduced profiling sensitivity due to the usage of outdated reference databases.

With these limitations in mind, there is a need for a tool optimized for monitoring with extensive features. To address this, we developed MMonitor, a software platform for real-time microbial monitoring based on ONT long-read sequencing platforms.

## DESIGN

We designed MMonitor considering the following:

- Intuitive user interfaces (graphical & command line) that appeal to researchers of varying levels of computational skill.
- Ability for time-series monitoring to track taxonomic shifts as new reads arrive.
- Customizable pipelines for 16S rRNA gene and WGS based taxonomic and functional analysis including metagenome assembly.
- Up-to-date databases with the ability to update automatically.
- Easy operation without previous installation
- Useable on high-end consumer grade hardware.
- Remote data access from any device or local processing.

A comparative analysis of our method with other tools in the real-time metagenomics space (e.g. minoTour [20], NanopoReaTA [21], MARTi [22], Nanometa Live [23], MAIRA [29], EPI2ME [30], BoardION [31], and CRuMPIT [32]) can be found in the Results section.

## RESULTS

### Metagenome monitoring software

Here we demonstrate MMonitor, an open-source software for automated monitoring through the real-time analysis of metagenomic nanopore sequencing data. The software consists of two components: a desktop application for running analyses and a web-based application for interactive result inspection. The desktop part integrates popular bioinformatics pipelines that users can run through a graphical user interface (GUI) or a command line interface (CLI), enabling analysis on local or remote systems. The method takes advantage of the real-time characteristics of nanopore data, allowing the automated analysis of metagenome reads as the sequencer continuously produces more data. The Web application features a visualization dashboard that provides users with dynamic insights into the taxonomy throughout time and the functional potential of metagenomes. It includes key metrics, such as quality scores, diversity indices, and taxonomy–metadata correlations, with options to export results for further analysis. We believe that MMonitor can reduce the time it takes researchers to address biological questions, making it useful in time-critical settings. It was developed and tested in collaboration with environmental biotechnologists, ensuring its usefulness for the tracking of metagenomes on site. Here, we describe how the software was designed to track the metagenome of multiple bioreactors over a month using 16S rRNA gene sequencing data. Then, we illustrate how MMonitor can use whole-genome sequencing (WGS) data to create annotated high-quality metagenome-assembled genomes (MAGs) for functional insights. Finally, we compare MMonitor to other existing tools in the area of metagenome monitoring.

Data that are sequenced at regular intervals (e.g., weekly samples in our study, though the frequency depends on the application) are automatically analyzed by MMonitor Desktop and the results are sent to the dashboard. The data is accessible locally or through a web browser. MMonitor offers two pipelines (Fig. **1** B), one for taxonomic analysis and one for functional analysis. For taxonomic analysis, it processes 16S rRNA gene sequencing and WGS reads, while the functional pipeline can only be run with WGS data. 16S rRNA gene sequencing provides quick taxonomic profiles and multiplexing allows analysis of more samples simultaneously, while WGS offers deeper taxonomic resolution and strain-level insights, but requires more time and computational resources. WGS enables *de-novo* assembly for full genome insights. We recommend the use of 16S rRNA gene sequencing for a general overview and WGS at key intervals for in-depth analysis. Refer to the methods section for pipeline details.

**FIG 1.**
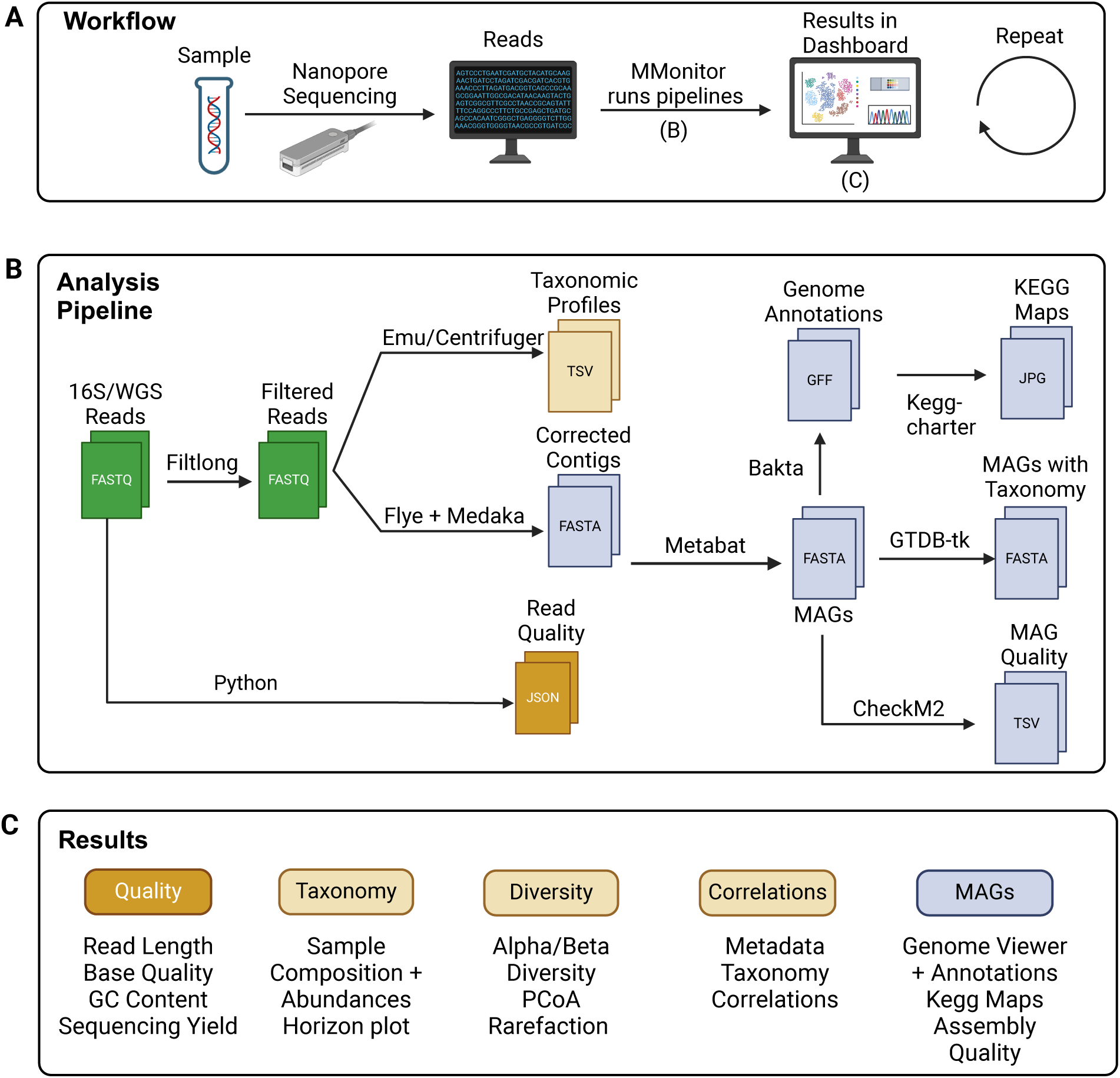
Workflow of MMonitor for metagenome tracking. (A) The general workflow of using the software in a laboratory. (B) Computational steps of the analysis pipeline. (C) The results available in the dashboard after analysis.

### 16S rRNA gene time series in three BES reactors

This section reports the nanopore 16S rRNA gene time series from three bioelectrochemical systems (BESR1–BESR3). We analyzed 66 amplicon libraries collected between July and August 2023 with MMonitor. Unless noted otherwise, “16S” refers to these BES datasets; genome-resolved WGS results from the AF and UASB reactors are presented separately.

The bioelectrochemical system (BES) originated from a parallel project that aimed to test whether removing H_2_ with electrodes could steer anaerobic fermentation pathways. The reactor achieved stable H_2_ removal and produced a 16S rRNA gene sequencing time series. We used these data primarily to validate MMonitor: frequent sampling under a defined perturbation provided an excellent test for real-time tracking, horizon plots, and diversity analyses.

Reads were quality-filtered using the 16S defaults (remove reads

*<* 1000 bp or

*>* 2000 bp, or with PHRED

*<* 10); the remaining reads were used for downstream taxonomic profiling and diversity analyses.

Here, we analyze the taxonomy of the three monitored bioreactors at different taxonomic ranks (Fig. **2**). Across the three BES reactors (R1, R2, and R3), the microbial communities were consistently dominated by members of the Bacillota phylum, followed by lesser contributions from Proteobacteria, Actinobacteria, and Bacteroidetes. During the time course examined, MMonitor revealed that the microbial composition (Fig. **2**A) of each bioreactor remains relatively stable in higher taxonomic ranks, with Bacillota continually dominating. However, at finer taxonomic resolutions, fluctuations were more noticeable.

**FIG 2.**
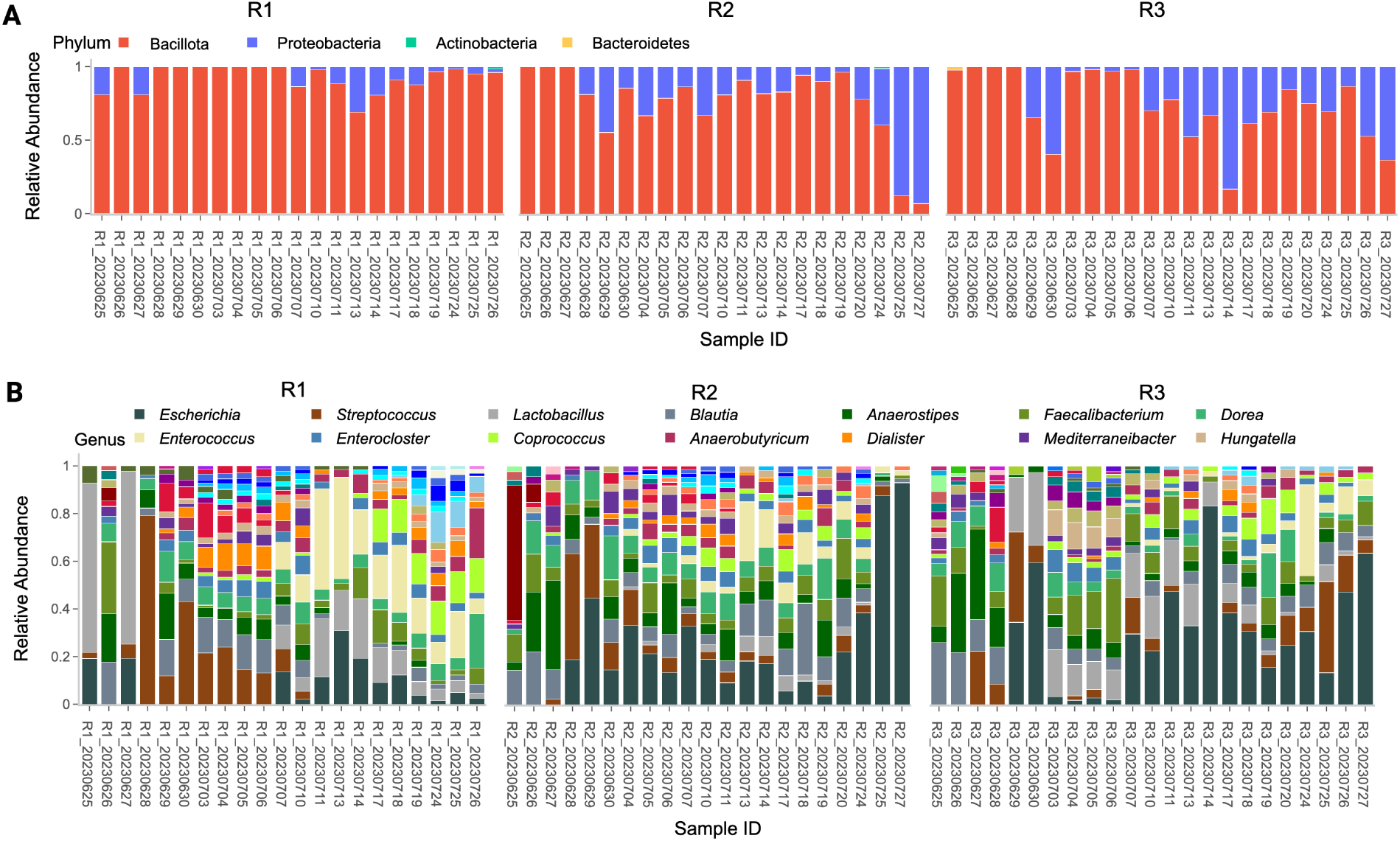
Taxonomic composition of the three bioreactors at different taxonomic ranks presented by MMonitor. For clarity, the legends only show the names of the 14 most abundant taxa. (A) Stacked bar plot of relative abundances of most abundant taxa found in the three bioreactors (R1, R2, and R3) at phylum level. (B) Stacked bar plot of relative abundances of most abundant taxa at genus level.

At the genus level, genera such as *Escherichia*, *Streptococcus*, *Lactobacillus*, *Blautia*, *Anaerostipes*, and *Faecalibacterium* dominated the bioreactor along with other minor genera (Fig. **2**B). In summary, there were some periodic variations, especially in finer taxonomic ranks, but at higher ranks, the community in all three bioreactors showed a more stable equilibrium in which Bacillota and Proteobacteria dominated the taxonomies.

The top 5 taxa in R1 at species level were *Enterococcus faecalis*, *Streptococcus salivarius*, *Escherichia coli*, *Faecalibacterium prausnitzii* and *Anaerostipes hadrus*. Although at the species level the taxonomic changes were more apparent, and while from day to day mostly minor changes appeared, throughout the weeks the taxonomy was unstable and certain taxa emerged while others disappeared (Fig. **3**A).

**FIG 3.**
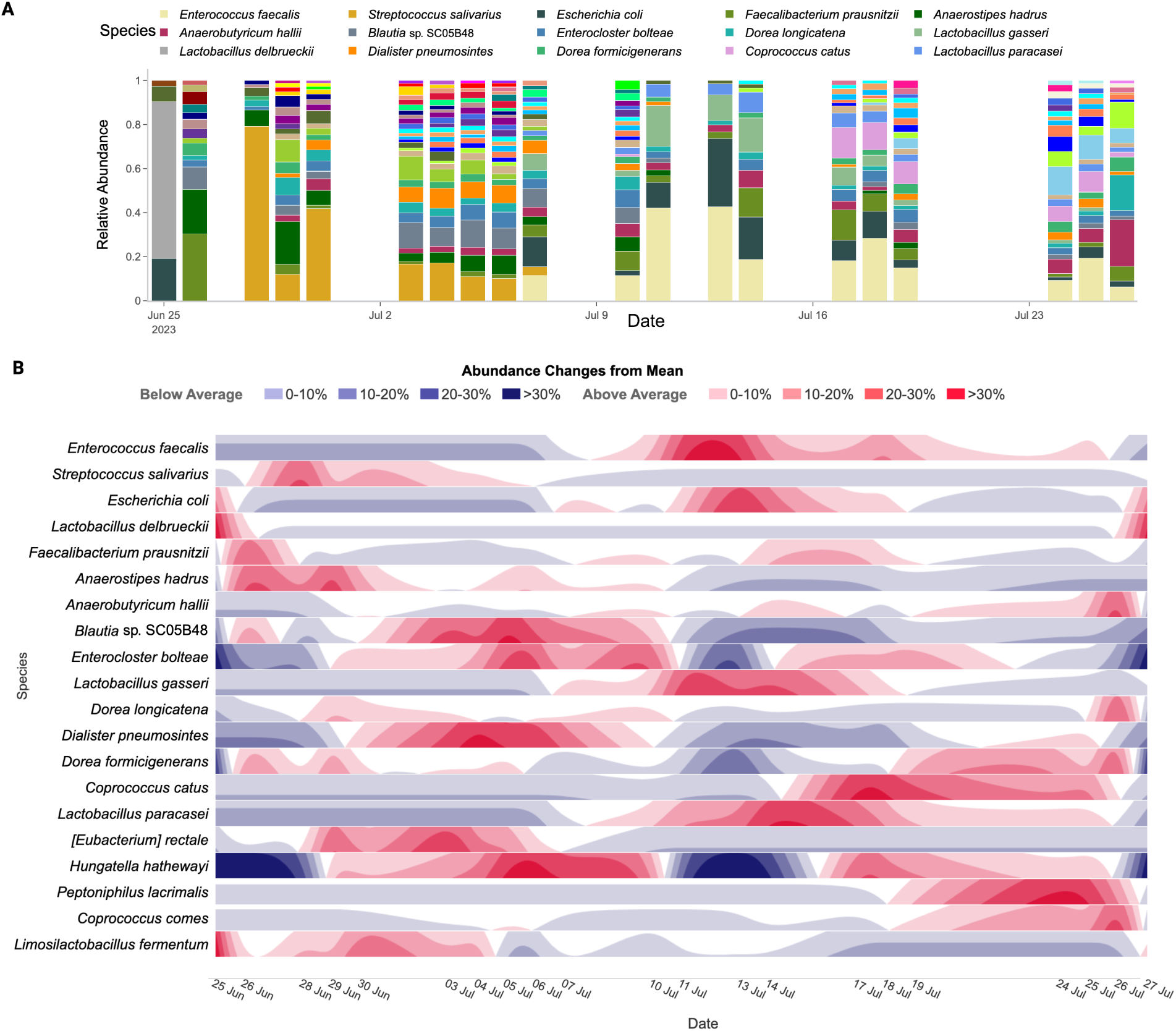
Taxonomic composition of BES reactor R1 presented by MMonitor. (A) Stacked bar plot of relative abundances of most abundant species found in R1 during the time of monitoring. (B) Horizon plot of the R1 taxonomy, visualizing changes in abundance over time.

These changes in abundance can be easily seen in a horizon plot (Fig. **3**B). It shows the changes in relative abundance of the top 20 species in R1 throughout time. Each row represents a species, red hills indicate an increase in abundance compared to the mean abundance, and blue bands indicate a decrease. At first, certain taxa, such as *S. salivarius* and *Dorea longicatena* increased in abundance, before decreasing again. In contrast, other taxa such as *E. coli* and *E. faecalis* did not appear until 10 July, resulting in a continuous blue band before that date, followed by increased abundances afterward, indicated by a red hill. In general, certain patterns become apparent, which could potentially indicate key time points. For example, around 11 July some event may have occurred that led to a sudden change and increase or decrease in overall abundance of many taxa. Similarly, on 25 June this can be seen for some taxa such as *Lactobacillus delbrueckii*, *F. prausnitzii* or *A. hadrus*.

Community shifts in the BES reactors were also described in detail in a previous dissertation based on the same dataset [33]. There, marked changes in community composition were observed at 201 h, 354 h, and 509 h, with differences between the three replicates and two enrichment phases in R1 (0–226 h and 251–559 h). These shifts were associated with changes in metabolite concentrations (e.g., acetate, propionate, *n*-valerate) and biofilm growth at the electrodes, which likely influenced hydrogen availability. However, the exact causes of the observed appearances and disappearances of species remain unresolved. As the goal of the present study was to demonstrate MMonitor’s ability to capture such dynamics in real time, we focused on showcasing the software’s monitoring capabilities rather than providing a full biological interpretation of these events.

Metadata such as the concentration of metabolite products can be correlated with the taxonomic abundance of taxa, possibly helping to understand their role in complex metagenomes. Additionally, MMonitor’s dashboard allows for the inspection of common alpha and beta diversity metrics, rarefaction curves, and clustering samples using principal coordinate analysis. The sequence quality metrics presented in the dashboard can help users filter low-quality samples and see when more strict filtering could be useful to obtain more significant analysis results.

### Analysis based on de-novo assembled MAGs and functional analysis

To demonstrate additional use cases of our software, we also applied it to WGS data. This is particularly important, as 16S rRNA gene sequences alone are often unable to differentiate between functionally distinct microbial taxa. Closely related species or strains with nearly identical 16S rRNA genes may possess different genomic content and functional capabilities [34]. WGS data also enables the tracking of other phylogenetic and functional marker genes beyond 16S rRNA, which is particularly valuable in complex biological environments where distinct taxa may fulfill similar ecological roles.

MMonitor successfully generated metagenome-assembled genomes (MAGs) and provided insight into the functional potential of individual microbes. From a nanopore WGS dataset, we obtained seven high-quality MAGs, which met the Minimum Information about a Metagenome-Assembled Genome (MIMAG) standards [35] and were assessed by CheckM2 [36].

Taxonomic annotation using GTDB-Tk revealed that six MAGs belonged to Bacteria and one to Archaea (Table **1**). Two MAGs were classified down to the species level: *Pseudoclavibacter_A caeni* and *Methanobacterium_C congolense*, with ANI values of 99.28% and 97.88%, respectively, indicating high sequence similarity to known reference genomes. The remaining MAGs were classified at the genus level, including *Dysosmobacter*, *Bulleidia*, and *Acetobacter*, suggesting they are novel species within these known genera. In particular, two MAGs (*JAEXAI01* and *JAUZPN01*) lacked close reference genomes and were assigned new placeholders at the genus level, indicating that they may represent novel genera (Table **1**). MMonitor also annotated the MAGs and mapped the annotations to the KEGG database to create metabolic maps that show the metabolic potential of a MAG. We have made the genomes, annotations, and metabolic maps of the MAGs available on GitHub. For details on accessing the data, please see the data availability section.

**TABLE 1.**
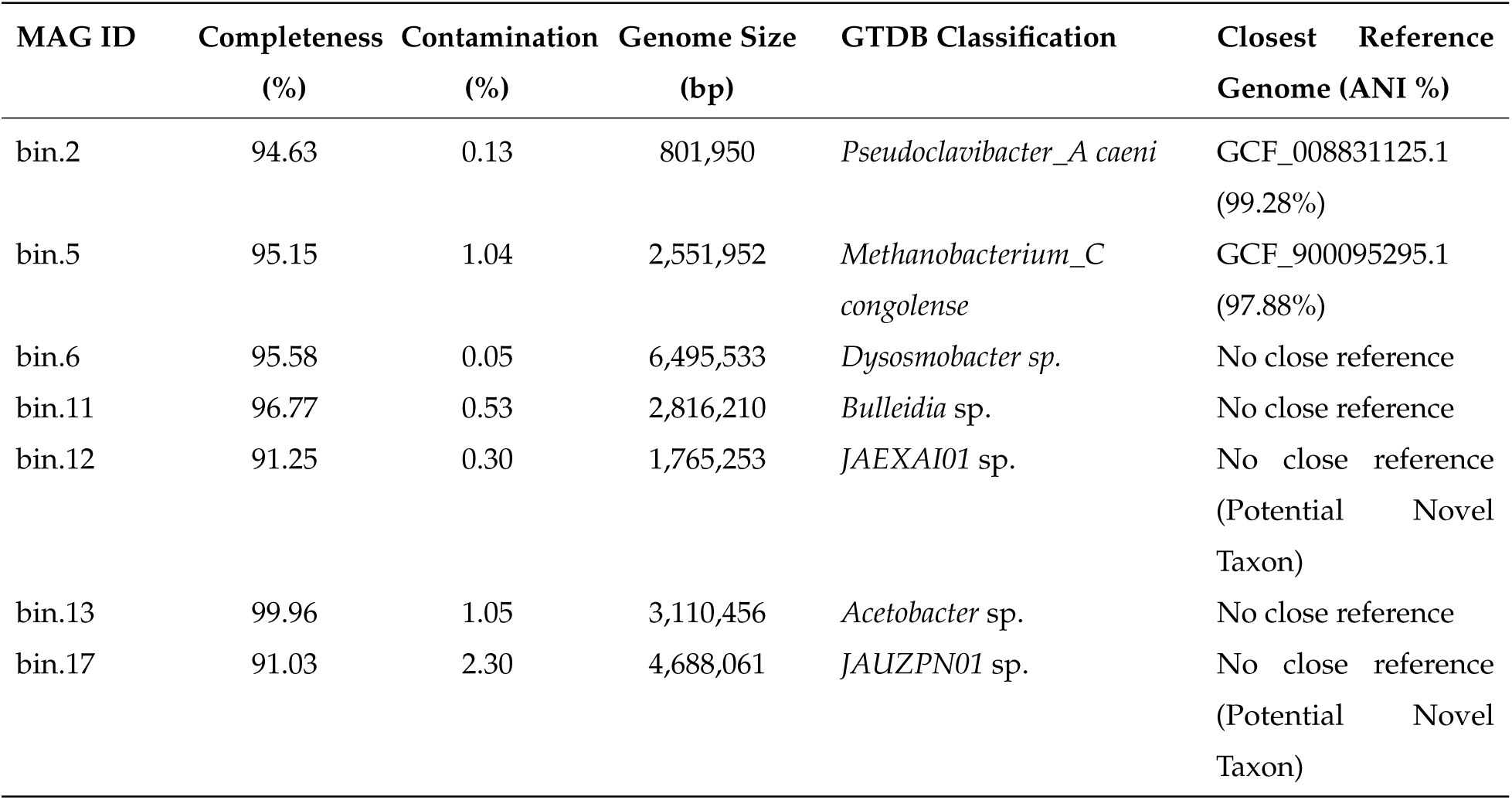
High-quality metagenome-assembled genomes (MAGs) with completeness ≥ 90%, contamination *<* 5%, genome size ≥ 100, 000 bp, and GTDB taxonomic annotation.

### Modified versions of popular algorithms for more efficient real-time monitoring

To optimize MMonitor for real-time tracking, we have made minor adjustments to taxonomic profilers without affecting result quality. Emu now optionally saves reference indices on the hard drive, avoiding recomputation during sequencing runs and improving runtime, especially with SSDs. For high-performance systems, users can increase the batch size when Emu uses minimap2, reducing input/output operations at the cost of more memory use. Centrifuger is modified to hold indices in memory and process new sequences immediately, lowering runtime for small batches where index loading is a bottleneck, enhancing efficiency in real-time scenarios. These changes do not alter the behavior of the core algorithm and can be disabled if necessary. We also introduced updated reference indices that can be downloaded from MMonitor for users who need current databases, along with an option to retrieve sequences and rebuild the indices of chosen domains.

### Feature comparison with similar software

To position MMonitor within the landscape of other metagenomics tools, we compared its features to those of existing software. The primary focus of this comparison was on tools that support real-time data processing, including minoTour [20], Nanometa Live [23], MARTi [22], NanopoReaTA [21], EPI2ME [30], boardION [31] and CRuMPIT [32] (see Table **2**). We examined these tools according to different criteria that influence the usability of a software for real-time monitoring of metagenomes.

**TABLE 2.**
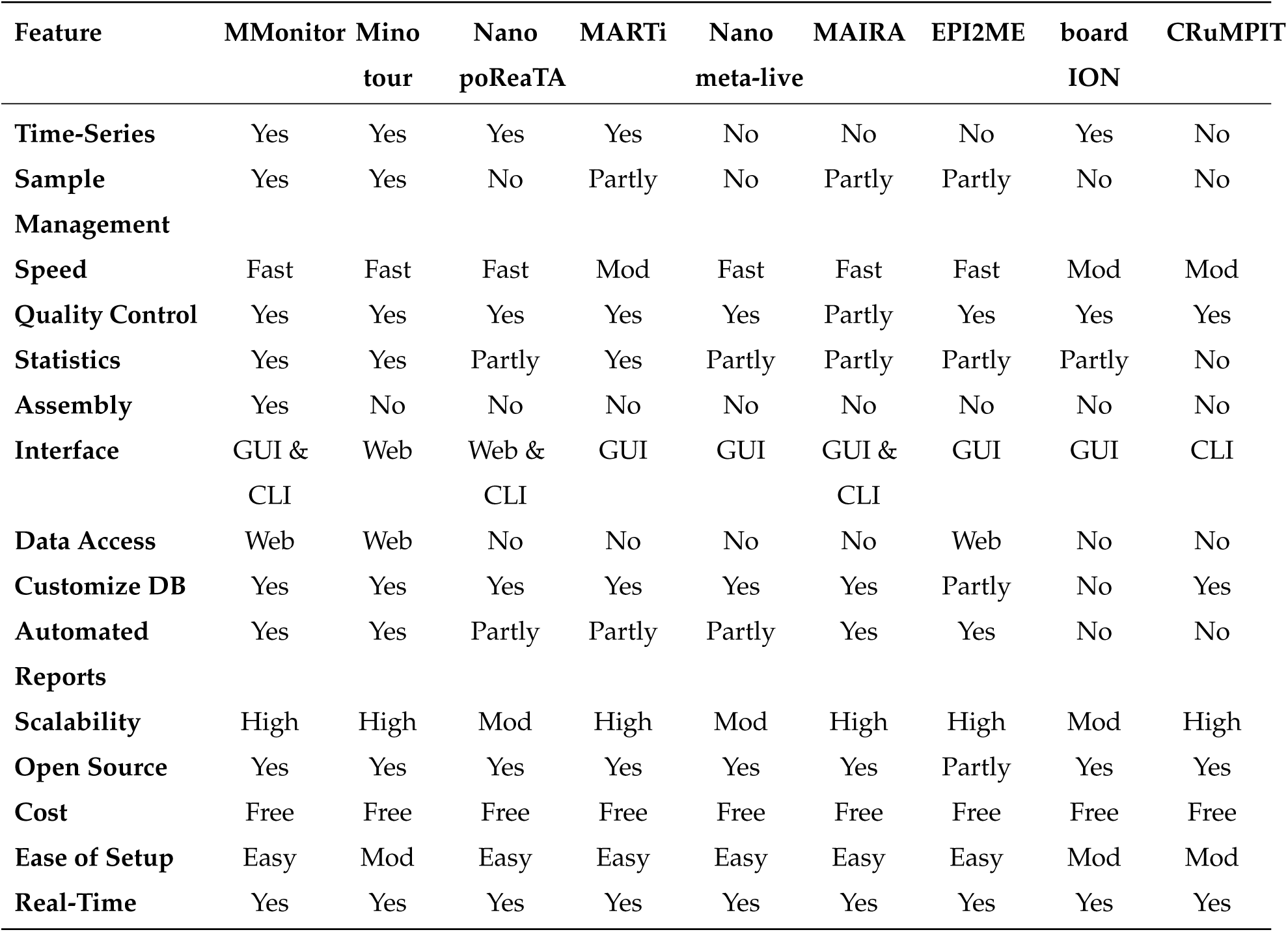
Comparison of eight real-time metagenome monitoring tools across 15 criteria. Criteria are scored as Yes (fully met), Partly, or No. Abbreviations: Mod = Moderate; GUI = Graphical User Interface; CLI = Command Line Interface. “No” under Data Access indicates local-only use, while “Web” denotes remote access.

| Feature | MMonitor | Mino<br>tour | Nano<br>poReaTA | MARTi | Nano<br>meta-live | MAIRA | EPI2ME | board<br>ION | CRuMPIT |
| --- | --- | --- | --- | --- | --- | --- | --- | --- | --- |
| Time-Series | Yes | Yes | Yes | Yes | No | No | No | Yes | No |
| Sample Management | Yes | Yes | No | Partly | No | Partly | Partly | No | No |
| Speed | Fast | Fast | Fast | Mod | Fast | Fast | Fast | Mod | Mod |
| Quality Control | Yes | Yes | Yes | Yes | Yes | Partly | Yes | Yes | Yes |
| Statistics | Yes | Yes | Partly | Yes | Partly | Partly | Partly | Partly | No |
| Assembly | Yes | No | No | No | No | No | No | No | No |
| Interface | GUI &<br>CLI | Web | Web &<br>CLI | GUI | GUI | GUI &<br>CLI | GUI | GUI | CLI |
| Data Access | Web | Web | No | No | No | No | Web | No | No |
| Customize DB | Yes | Yes | Yes | Yes | Yes | Yes | Partly | No | Yes |
| Automated Reports | Yes | Yes | Partly | Partly | Partly | Yes | Yes | No | No |
| Scalability | High | High | Mod | High | Mod | High | High | Mod | High |
| Open Source | Yes | Yes | Yes | Yes | Yes | Yes | Partly | Yes | Yes |
| Cost | Free | Free | Free | Free | Free | Free | Free | Free | Free |
| Ease of Setup | Easy | Mod | Easy | Easy | Easy | Easy | Easy | Mod | Mod |
| Real-Time | Yes | Yes | Yes | Yes | Yes | Yes | Yes | Yes | Yes |

### Detailed feature comparison

All tools in table **2** support real-time processing of nanopore data, but implement alternative algorithms and report different downstream analyses. For example, MinoTour, MARTi, NanopoReaTA, and Nanometa Live offer pipelines for real-time taxonomic identification, but do not support *de-novo* metagenome assembly. Of the compared tools, only MMonitor implements a complete metagenome assembly pipeline with the ability to generate MAGs from long-read data only. EPI2ME includes an assembly pipeline for single isolates, but not for community-level metagenomes.

NanopoReaTA is designed specifically for differential transcriptome data and gene-specific analyses. It is useful for taxonomic analysis but lacks the capability to integrate different types of data and methods besides cDNA, which limits its utility for monitoring microbiomes. It includes a Docker container that can be used to run the software. While this ensures portability, it also requires command-line usage to operate and will not work without Docker. This could be a drawback if used on a system without root access. Also, it is limited to usage on Linux based systems. Despite these drawbacks, its ability to perform real-time cDNA analysis makes it a useful asset for monitoring transcription, complementing other tools that do not support this.

MARTi is a tool for clinical and research-focused metagenomics similar to MMonitor. It offers real-time pathogen detection and some taxonomic classification capabilities and reports antimicrobial resistance genes, but lacks time-series plots and correlation analysis. Although it offers rarefaction analysis, it does not report other diversity metrics. It has an excellent GUI to run pipelines locally or remotely on a server but requires extensive setup of a back-end engine and the front-end using the command line. In conclusion, MARTi is a capable tool for real-time monitoring of metagenomes, especially if time-series analysis is not the main focus.

Nanometa Live is another tool designed for the real-time analysis of metagenomic data from ONT devices. It integrates Kraken2 [37] for taxonomic profiling and provides a user-friendly interface to visualize data as they are generated. It can operate offline, making it a viable solution for fieldwork. However, it has limited capacity for more in-depth functional analysis, as it lacks methods for genome assembly or annotation. Its focus is primarily on identifying species, rather than exploring functional pathways or gene-level insights. Although it offers multiple ways to install, including a Docker container, the lack of pre-built binaries requires the use of command line interfaces for installation and operation. Its execution is restricted to running locally and uses Kraken2 for profiling, which requires more RAM than Centrifuge, possibly problematic for standard lab computers. Although the tool performs a sequence quality assessment, it does not offer other downstream analyses. In summary, Nanometa Live addresses some issues that make many tools unattractive and not accessible to a broader audience.

However, further functionalities for monitoring metagenomes such as diversity and correlation analysis, or time-series plots, are lacking, and its operation can be hard for the average user with its CLI dependency for operation and installation.

MAIRA (Mobile Analysis of Long Reads) is an open-source software designed for the real-time taxonomic and functional analysis of long-read metagenomic sequencing data on a laptop. MAIRA focuses on real-time analysis of sequencing data as it is generated, but lacks support for time-series visualizations or analyses involving multiple time points; it does not provide tools for tracking changes throughout time or managing multiple samples during extended periods. Optimized for high-end laptops (≥ 32 Gb of memory, 500G of SSD), it performs fast genus-level analysis and on-demand species-level analysis, making it suitable for field use with limited computational resources, but does not include sequence quality control features, advanced statistical analyses, or diversity metrics. Focusing on taxonomic classification and functional annotation, including antibiotic resistance and virulence factors, MAIRA offers a GUI and CLI mode but lacks remote data access. It is open-source, free, and easy to install on all major OS due to its Java base. Although useful for real-time analysis, MAIRA lacks time-series visualizations and sample management for long-term monitoring.

EPI2ME, by Oxford Nanopore Technologies, is a cloud-based platform for real-time taxonomic analysis using WGS or 16S rRNA gene sequencing data. It includes a genome assembly and variant calling pipeline for isolates, but not metagenomes. Specialized workflows are available, such as WIMP analysis for 16S rRNA gene sequencing data and metagenomics with WGS data. Results can be accessed locally or from the cloud, though only via the EPI2ME-labs desktop software, not a browser. Users cannot customize workflows without modifying code, and using Nextflow requires configuration knowledge or pre-made files. Although EPI2ME provides Open Source pipelines, the software itself is not Open Source. It handles large single samples well but lacks multi-sample processing without multiplexing. Free for customers, EPI2ME is user-friendly with a strong GUI and supports Windows through containerization.

However, it does not support custom databases, multi-sample or time-series analyses, nor does it offer species-level resolution for metagenome monitoring. Although comprehensive plots for taxonomy are included, advanced analysis like PCoA requires manual data export. The absence of metagenome assembly limits it to metagenomic taxonomic analysis.

MinoTour is a tool for real-time monitoring of nanopore sequencing runs, allowing users to track sequencing progress, report sequencer performance, and perform real-time taxonomic profiling. It integrates pipelines for alignment, metagenomics, and adaptive sequencing, and supports custom workflows such as the ARTIC pipeline for viral genomes like SARS-CoV-2. While its Django-based architecture provides extensibility, MinoTour lacks downstream metagenomic analysis and advanced statistical tools, limiting its application for broader microbiome studies. In addition, its web interface is convenient but offers limited customization options, and the requirement for users to host their own instance makes it less accessible to non-technical users.

BoardION, while not directly aimed at metagenomic analysis, focuses on monitoring the sequencing process in real time. This tool is highly useful for tracking sequencing progress and instrument health, but lacks functionality for taxonomic profiling, genome assembly, or functional analysis. Its use is limited to the sequencing process, rather than downstream analysis of microbial communities.

CRuMPIT is a software solution developed for clinical metagenomics, particularly with a focus on pathogen detection. Like other real-time tools, it integrates workflows and processes data quickly for pathogen identification. However, it lacks extensive statistical analysis and the broader community ecology features that MMonitor and similar tools offer. Its primary strength lies in its ability to process and analyze clinical samples for pathogen surveillance, which is highly useful in medical and clinical environments but may not be as relevant for more diverse ecological or industrial applications.

MMonitor was designed to meet all the criteria that are missing in the other tools. The prebuilt binaries require minimal setup, its code is open-source, it combines a straightforward GUI with a CLI, and supports remote computation, making it accessible to many users. It enables real-time analysis, crucial for experimental adjustments, and features databases that can be seamlessly modified or updated directly from the GUI. The system helps to organize samples, enabling simple sample comparisons. MMonitor also provides custom visualizations, such as the horizon plot, allowing researchers to quickly recognize trends. Analyses and result evaluations can be conducted locally or remotely, which is beneficial when internet access is limited or data privacy is a concern. The MMonitor pipelines offer sufficient scalability for handling numerous and large samples, and its dashboard visualizations help with data analysis regardless of sample size. We built the software for multiple platforms, together with all dependencies, making it portable and enabling installation and usage without requiring command-line knowledge. In a similar manner, we created a Docker container for the Dashboard, enabling researchers to easily host their own version if they require greater data control.

Additionally, MMonitor Desktop offers an offline mode that can initiate a local dashboard to inspect the results. MMonitor delivers metagenome monitoring features that surpass existing tools, effectively addressing limitations of other software.

In addition to these direct competitors, we conducted a supplementary comparison with broader metagenomic analysis tools such as MG-RAST, MEGAN, QIIME 2, and MetaPhlAn 4 (see Table **3** in the supplementary material). Although these tools offer powerful taxonomic and functional profiling, they generally lack the real-time capabilities necessary for monitoring dynamic environments.

**TABLE 3.**
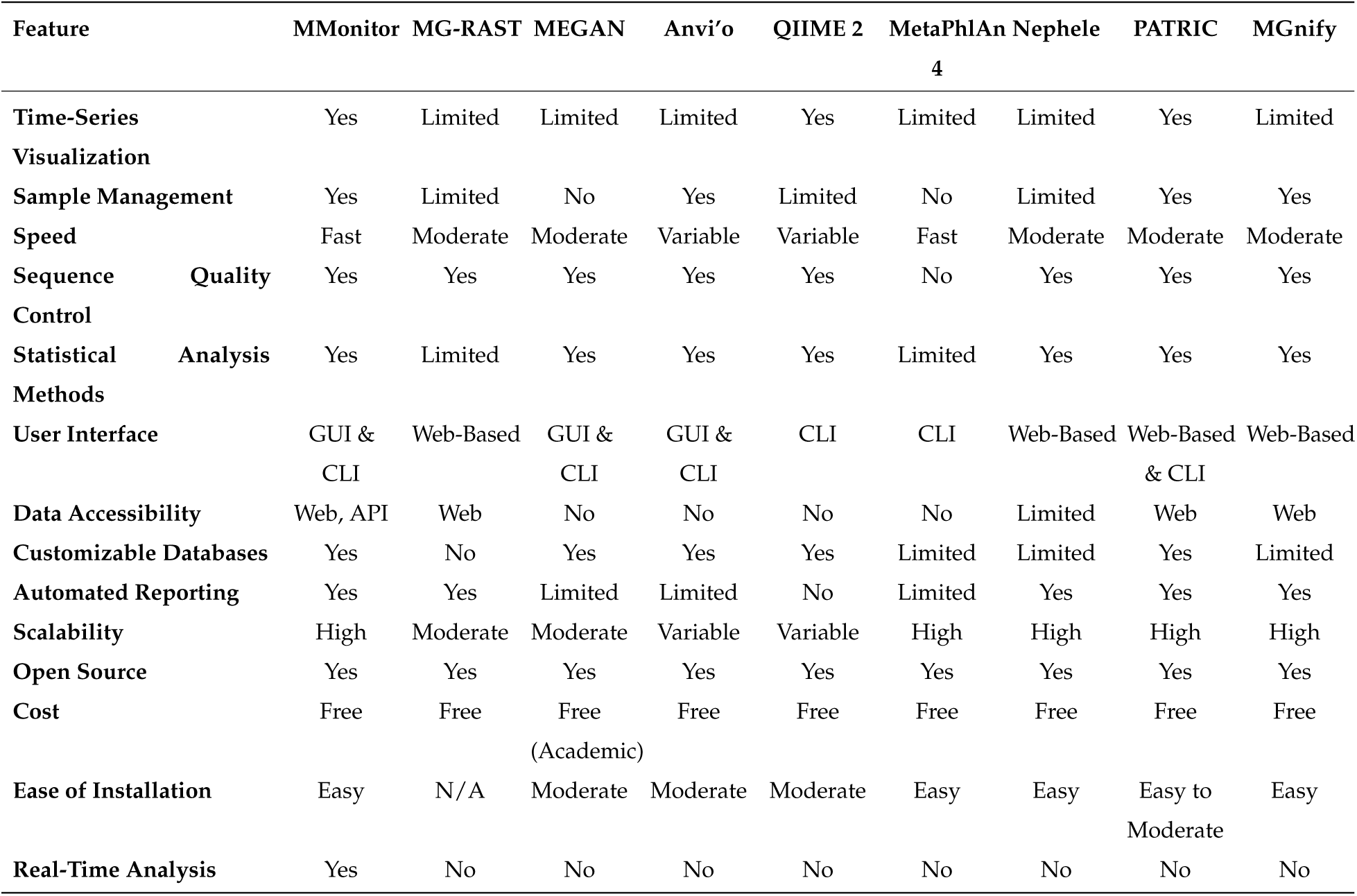
Comparison of MMonitor with Indirect Competitors (Broader Metagenome Analysis Tools). N/A: Not applicable.

## DISCUSSION

Recent advances in sequencing technologies, along with the growing number of studies applying these methods in clinical, environmental, and industrial microbiology, reflect the increasing interest in metagenome monitoring and highlight the need for accessible and optimized tools for this purpose. However, the costs associated with metagenome monitoring, including sequencing expenses (at least $13 per sample for 16S rRNA gene sequencing and $50 per sample for WGS sequencing with multiplexing) and computational resources, can be substantial. Although more expensive than short-read sequencing, Nanopore sequencing enables species- and even strain-level resolution and real-time analysis, but with a higher error rate. Real-time local analysis reduces infrastructure needs and allows on-site sequencing, but consumables such as flow cells, reagents, and computational capacity must also be considered. Alternatives to WGS-based monitoring include targeted molecular assays, such as hybridization probe arrays. These approaches can quantify specific microbes with high sensitivity; however, they are restricted to known targets and therefore cannot capture the overall composition of the community.

### Possible applications for monitoring

MMonitor has a diverse range of applications, not limited to certain environments or species. Instead, it can be used for any nanopore sequencing data, either from metagenomes or isolates, and for whole genome sequencing or targeted sequencing. In environmental biotechnology, it can be used to monitor microbial populations in bioreactors, which can aid in the optimization of production processes for biofuels or bioremediation efforts.

Metagenomic monitoring can uncover novel functional genes and enzymes, including those from uncultivable microorganisms, that can be harnessed to improve biomass conversion and biofuel production [38]. It can also track shifts in microbial community composition and activity, providing the information needed to optimize and control microbiome behaviour for greater yield, stability, and a broader range of products in large-scale biomanufacturing systems, as emphasized by the advantages and process stability requirements mentioned in [39]. In clinical microbiology, real-time monitoring of microbial communities can assist in tracking pathogen emergence and antibiotic resistance patterns. The ability to correlate microbial abundances with environmental or clinical metadata enhances the utility of MMonitor in these contexts. Although metagenome monitoring has not yet been standardized as part of routine production workflows, several large-scale plant-level applications demonstrate its practical feasibility. For example, shotgun metagenomics has been used to characterize resistomes in wastewater treatment plants [40], to profile microbial community dynamics across sewer networks [41], and to monitor bioleaching microbial communities via high-throughput sequencing [42]. These studies illustrate that, although sequencing turnaround currently limits fully real-time feedback, metagenome monitoring is already integrated into industrial workflows and is expected to expand as sequencing costs and analysis latency decrease.

### Relevance of reactor systems to this study

MMonitor was validated in two complementary contexts. BES reactors provided short, controlled enrichments with frequent 16S sampling, enabling rapid taxonomic tracking and diversity analyses. In contrast, AF and UASB reactors from a separate MCFA project supplied long-running WGS datasets to demonstrate genome assembly, binning, and functional monitoring.

Together, these systems highlight the dual use cases of MMonitor: (i) rapid 16S-based tracking (BES) and (ii) genome-resolved functional monitoring in continuous bioprocesses (AF/UASB). The BES itself was only indirectly related to MMonitor, but its time series offered a stringent test case for our 16S pipeline.

### Limitations and Future Development

Despite its strengths, MMonitor has certain constraints and areas for future improvement.

- High sequencing loads: Very frequent data updates (e.g. when running more than one sequencer) may overburden consumer-grade hardware, as current optimizations focus on moderate analysis intervals.
- OS restrictions: Although the desktop app runs on multiple platforms, the underlying bioinformatics pipelines currently require a Unix-based environment. Full native support for Windows is not yet available.
- Short-read integration: MMonitor is presently optimized for nanopore data. The WGS pipeline uses a database built on reference genomes, which works on both long- and short-read data (e.g., Illumina), but using MMonitor with short-read data has not been extensively tested. Full short-read support, including a short-read 16S rRNA gene pipeline and an assembly pipeline, is planned for future releases. Other

long-read technologies should work out of the box, but have not yet been tested.

- Database Limitations: Default references focus on bacteria, archaea, and fungi. Viral or other specialized targets may require manual database creation and indexing.

We intend to address these limitations in future releases by working on the following features.

- Containerization: Using tools like Docker or Singularity to containerize the pipelines could improve cross-platform compatibility, allowing them to run on Windows and other operating systems more easily.
- More functional analyses: Monitoring antibiotic resistance genes and incorporating methylation detection (via Dorado) could be useful for pathogen surveillance and microbial regulatory studies.
- Support for more input data: Integrating transcriptomic, proteomic, or metabolomic data and support for sequencing data from other technologies could allow more advanced analyses and increase the range of applications.
- Performance optimization: GPU-accelerated algorithms and more efficient I/O handling and further improvements to the databases could improve performance and help with growing data sizes.

Another promising direction is the automation of sampling and sample preparation. Manual collection, DNA extraction, and library preparation remain time-consuming and can introduce variability. Many wastewater treatment facilities already deploy automated autosamplers as part of routine monitoring workflows [43], and open-source automation platforms (e.g., Opentrons and VolTRAX) are emerging to standardize metagenomic library preparation protocols [44].

MMonitor is an open-source project hosted on GitHub, and we welcome contributions from the community. We encourage anyone interested in reporting issues, suggesting new features, or contributing code to participate in its ongoing development.

### Conclusion

In summary, MMonitor addresses current limitations in real-time metagenome analysis by providing a fully integrated solution that combines rapid taxonomic profiling, de novo assembly, and intuitive time-series visualization. By automating data analysis and many other key steps, such as updating reference databases and sample management, MMonitor enables users to capture dynamic shifts in microbial populations. Our validation on bioreactor samples demonstrated its reliability for both 16S rRNA gene and WGS data, and the comparison with existing tools highlights the strengths of different approaches. As an open-source project, MMonitor welcomes contributions from the scientific community, fostering continuous improvement and broader adoption across diverse microbiome research domains. Looking ahead, the integration of automated sampling and sample preparation workflows could further enhance the utility of MMonitor by enabling near-continuous metagenome monitoring in industrial and clinical settings.

## STAR Methods

### Resource availability

#### Lead contact

Further information and requests for resources, data and software can be directed to Timo N. Lucas.

#### Materials availability

This study did not generate new unique reagents. All software scripts are distributed with MMonitor and available via the project repository. Reactor designs and cultivation protocols are described in detail in this study and referenced publications.

#### Data and code availability

- Raw 16S rRNA gene sequencing and WGS data are deposited at NCBI under BioProject **PRJNA1216071** (https://www.ncbi.nlm.nih.gov/bioproject/PRJNA1216071).
- MMonitor source code, built binaries, and usage instructions: https://github.com/lucast122/mmonitor.
- Analysis files created for this manuscript and the manuscript source: https://github.com/lucast122/mmonitor-paper.
- Database-building scripts are included in the MMonitor repository. Required databases can be downloaded using the graphical user interface of MMonitor. They are also referenced in the GitHub repository.
- A hosted instance of the MMonitor dashboard is available at https://www.mmonitor.org.

### Experimental model and subject details

#### Microbial samples

Two distinct reactor systems were studied. For 16S rRNA gene sequencing experiments, three bioelectrochemical system (BES) reactors were inoculated with fresh human fecal samples collected with informed consent and processed anaerobically (see Method details). For WGS experiments, two continuous bioreactors (anaerobic filter and UASB) were inoculated with a chain-elongating microbiome derived from a previously operated CSTR and maintained over a period of ∼1000 days. This experiment was approved by the Ethics Committee of the Medical Faculty of the University of Tübingen (Project number: 456/2023A). Anonymized stool samples were collected with the permission of all human subjects.

### Method details

#### Analysis pipeline

MMonitor was written in Python (v3.11). The desktop application handles data input and running pipelines, the computations themselves are performed by external bioinformatics tools (Fig **1**B), for which benchmarks and methods can be found in the respective publications. All input reads are first passed to Filtlong (v0.2.1) for initial quality filtering. Filtering parameters can be defined in the Analysis Configuration tab of the MMonitor Desktop application, by default WGS reads shorter than 2000bp and 16S rRNA gene reads shorter than 1000bp or longer than 2000bp are discarded, as well as reads with PHRED score below 10. Only reads passing quality filtering are used for the remaining steps. Taxonomic analysis using nanopore WGS reads is performed by Centrifuger [45], which is based on the Burrows-Wheeler transformation (BWT) and the Ferragina-Manzini (FM) index [46, 47]. Centrifuger works similar to Centrifuge [48], but uses a run-block compression algorithm to reduce the memory footprint while only slightly increasing runtime. For nanopore 16S rRNA gene taxonomic profiling, MMonitor uses Emu [12], which uses minimap2 [49] for alignment and an expectation maximization (EM) algorithm [50] to refine relative abundances. Basic quality statistics for input reads are computed directly from input reads for quality score, read lengths, and number of bases using biopython seqIO [51]. For functional analysis, MMonitor requires WGS reads as input. The assembly pipeline was inspired by previous work [52]. Flye [53] is used to assemble the metagenomic reads into contigs. By default, the –meta parameter is used and –nano-raw is used to input reads. If required by the user, –nano-hq can be used instead if basecalling was performed using a sup model. The assembled contigs are corrected using medaka (v2.0.1) [54]. By default medaka tries to select the model automatically, but this is not always possible, for example if the data was called with an old basecaller or is not in pod5 format. In that case, the user needs to provide the correct model that fits the sequencing data. Detailed information on this can be found at https://github.com/nanoporetech/medaka. The resulting consensus assembly is binned using Metabat2. The resulting bins are taxonomically annotated by GTDB-tk (v2.4.0) [55]. The completeness and contamination of the resulting MAGs are then controlled using CheckM2 [36] and all MAGs with completeness above 80% and contamination below 10% are sent to the database. Bakta (v1.9.4) [56] is used to create annotations and then those are mapped to the KEGG database using keggcharter (v1.0.2) [57, 58]. In addition to the graphical interface, MMonitor can be run in command-line mode, enabling remote execution on other systems (e.g., via SSH) as described in the project README.

#### Webserver and Dashboard

The web server backend was designed with the Django framework [59] and the dashboard was implemented using a combination of Plotly Dash and Javascript [60]. Docker was used to bundle the dependencies of the server and can be used to deploy it. We offer a default online version of the MMonitor server that all users can access. However, if you set up your own MMonitor server, you gain full control over the database and user management, allowing you to customize access and data handling according to your needs.

In addition to visualizations, the dashboard also provides several statistical methods. Normalized counts *c_n_* (1) are calculated from raw counts *c_r_*, the number of aligned bases b, and a scaling factor f.

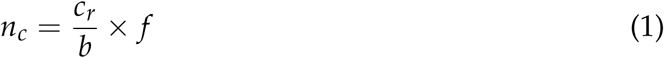

The scaling factor is the average number of aligned bases in all samples. It ensures that the raw and normalized counts have a similar magnitude. Alpha and beta diversity are calculated using scikit-bio diversity (v0.5.9) using normalized counts on demand. For alpha diversity, the Shannon (2) and Simpson (3) indices can be used, where *s* is the number of unique taxa and *p* is the proportion of the community represented by taxon *i*.

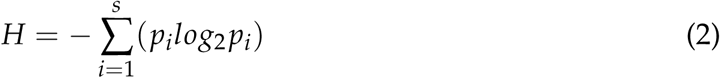

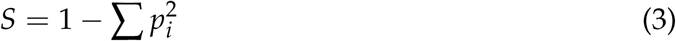

Beta diversity is determined by calculating the Bray-Curtis (4) distance between all samples, where *u_i_* and *v_i_* are the proportions of taxon *i* in sample *u* and *v*.

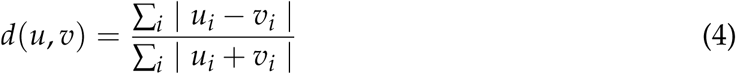

Principal coordinate analysis (PCoA) is performed using scikit-bio stats ordination. Horizon plots were implemented using the JavaScript D3 plotting library for each taxon, displaying the difference in abundance from the previous sample. Pandas was used to calculate taxonomy-metadata correlations using either one of the Pearson (5), Kendall (6), or Spearman (7) correlation coefficients. Here, *cov* is the covariance, *X* is a series of metadata values, *Y* is the series of taxon counts, *C* and *D* are the number of concordant and discordant pairs and *R(X)* is a function that returns the rank of the variable *X* [61, 62].

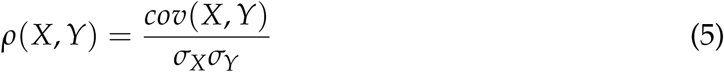

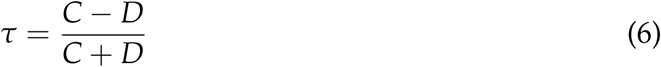

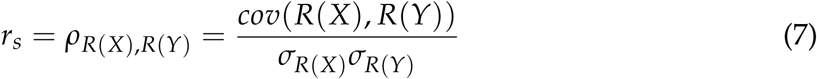

#### Reference database creation

MMonitor utilizes customized databases for taxonomic classification. The Database Manager window can be used to change profiler indices or to download and build fresh versions on demand. To save build time for users, we provide pre-built indices online (see data availability section). To create the Centrifuger index, we downloaded all complete genomes of bacteria, archaea, and fungi from the NCBI RefSeq database [63] (as of October 2024) and concatenated them into a single file. We also downloaded the taxonomy tree and taxonomy mapping file from the NCBI taxonomy dump. The centrifuger index was then built using the centrifuger-build command with default parameters, providing the concatenated genomes, taxonomy tree, and mapping file previously downloaded.

For the Emu database we followed the authors’ in their methods section Emu 16S database. All 16S rRNA gene sequences for bacteria and archaea from the NCBI targeted loci directory and the rrnDB-5.9 sequences were concatenated into a single file. Then we downloaded the NCBI taxdump and accesion-to-taxid mapping file and the mapping file for rrnDB-5.9. We used Emu’s build-database and Centrifuger’s internal script for building the databases. We did not include the index files in the MMonitor binaries due to their large file size and instead uploaded them to a public repository. If no index is found or provided by the user during run-time, MMonitor will notify the user and try to download it. The scripts for downloading and building the new databases are included in MMonitor and can be run on demand from the Database Manager of the desktop app. You can find the source code and custom index files in the Data Availability section.

While the 16S rRNA and WGS pipelines in MMonitor use different tools and databases, the WGS database is built from complete reference genomes (by default all RefSeq bacterial, archaeal, and fungal genomes). This design enables detection of any marker genes present in these genomes, making the same database creation scripts applicable to marker gene sequencing data, provided the target genes are part of the reference genomes. In future versions, we plan to add an option for users to supply a FASTA file with specific marker gene sequences, allowing the creation of dedicated marker-gene databases for faster querying and targeted monitoring.

#### 16S rRNA gene nanopore sequencing

For 16S rRNA gene amplicon sequencing, we used the 16S rRNA gene Barcoding kit 1-24 (SQK-16S024, Oxford Nanopore Technologies) and an R9.4.1 flow cell (FLO-MIN106). The 16S rRNA gene Barcoding kit enables rapid and full-length 16S rRNA gene sequencing for organism identification by using universal primers 27F (5′− AGAGTTTGATCMTGGCTCAG −′) and 1492R (5′-CGGTTACCTTGTTACGACTT −3′). For 16S rRNA gene amplification via PCR, we mixed 10*µ*l of 10ng DNA with 25*µ*l of LongAmp^(^*^R^*^)^ Hot Start Taq 2x Master Mix (New England Biolabs),10 *µ*l of an individual 16S rRNA gene barcode and 5*µ*l of nuclease-free water. The PCR cycles were 1 min 95°C, 25 cycles of 20s 95°C, 30s 51°C, 2min 65°C and a final elongation of 5min at 65°C. Afterward, bead cleaning, preparation of the cleaned up DNA library with rapid adapter and pooling, and preparation and loading of the R9.4.1 flow cell were performed following the manufacturer’s protocol. The sequencing time depended on the number of barcodes pooled in one library. We decided to sequence for 2h per barcode, resulting in a reasonable coverage of the 16S rRNA gene for better classification results. MINKNOW software version 23.11.4 was used for data generation.

#### Bioelectrochemical system setup (16S rRNA gene sequencing experiments)

The BES consists of two glass chambers (working and counter) separated by a Nafion 117 ion-exchange membrane (Sigma), so that the human fecal sample contacts only the working electrode *E_we_*. A three-electrode configuration (*E_we_*, *E_ce_*, *E_re f_*) was used with the working electrode poised at +350 mV vs. Ag/AgCl for H_2_ removal by oxidation. Both *E_we_* and *E_ce_*were carbon cloth electrodes (area 122 cm^2^). *E_we_* was spray-coated on both sides with platinated carbon (10%) at a loading of 4 mg cm^−2^. The reference electrode *E_re f_* was Ag/AgCl (silver wire chloridized, embedded in 3 M KCl and 0.7 g agarose).

Prior to assembly, Nafion 117 membranes were activated in 1 M sulfuric acid for 24 h and equilibrated for 24 h in a salt solution (85.5 mM NaCl, 16.9 mM Na_2_HPO_4_·2H_2_O, 41.9 mM NaHCO_3_). Reactors (including the membrane-separated electrodes) were assembled in a sandwich configuration and sterilized at 121^◦^C for 20 min under aerobic conditions. After autoclaving, chambers were filled with anaerobic, reduced medium and sparged with N_2_:CO_2_ prior to inoculation. Working volume was 21 mL in both chambers. The geometry minimized the microbe–electrode distance to ∼1 mm.

#### DNA extraction for BES (16S)

For DNA extraction, 500 *µ*L BES liquid was centrifuged (19,000×*g*, 5 min, 4^◦^C) to pellet cells. DNA was isolated with the AllPrep^®^ PowerFecal Pro DNA/RNA kit (QIAGEN) using a FastPrep-24^TM^ 5G bead-beater (2 cycles at 6.0 m s^−1^ for 40 s with 30 s rest). We followed the manufacturer’s instructions with one exception: < 450 *µ*L (instead of 300 *µ*L) supernatant was taken in pretreatment step 6 (after adding CD2), following the vendor’s guidance for higher volumes. DNA quantity was measured with Qubit^TM^ Flex (Invitrogen) and quality with NanoPhotometer^TM^ N60/N50 (Implen).

#### Continuous bioreactors with pertraction system (WGS experiments)

Two independently operated, continuously fed upflow bioreactors with integrated liquid–liquid extraction (pertraction) were run in parallel for ∼1019 days. Each bioreactor was a double-walled glass column (height 125 cm) maintained at 30^◦^C and pH 5.5. One vessel contained wheel-shaped plastic packing (anaerobic filter, AF reactor); the second was an upflow anaerobic sludge blanket (UASB) reactor without packing to promote granule formation. Fermentation broth was recirculated through two hollow-fiber membrane contactors: medium-chain carboxylates were extracted into a hydrophobic solvent and subsequently transferred into an alkaline extraction solution. An in-line 0.8 L filter module protected the membranes during early operation.

*Medium and inoculum:* both reactors were inoculated with 2 mL glycerol stock (0.04% v/v) from a chain-elongating microbiome previously maintained in a CSTR on ethanol/acetate feed. The defined medium contained ethanol (13.42 g L^−1^; ∼600 mM C) and acetate (3.12 g L^−1^; ∼100 mM C) at a 6:1 molar ratio, plus bicarbonate and nutrients.

*Operating periods:* over 1019 days, four periods (I–IV) were defined by construction and oxygen-management changes: limiting oxygen intrusion (airlocks, N_2_ sparging, metal tubing), adjusting reducing agents (L-cysteine, sodium sulfide), and later controlled O_2_ supply. Gas additions included N_2_/H_2_ (95/5% v/v; 0.3 ± 0.1 mL h^−1^) and air (11.8–17.1 mL h^−1^) during late operation. These interventions altered redox balance and *n*-caprylate productivity.

*Sampling:* biomass for metagenomics was collected from reactor columns and the filter module at Days 9, 283, 397/455, and 701.

### Quantification and statistical analysis

The statistical procedures used by the dashboard are defined in the Webserver and Dashboard section (Eqs. 1–7). Briefly, normalized counts are computed as 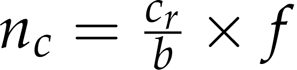; alpha diversity is estimated by Shannon and Simpson indices; beta diversity by Bray–Curtis distances followed by PCoA; and taxonomy–metadata associations are evaluated with Pearson, Kendall, or Spearman correlation using pandas and scikit-bio (v0.5.9). Where applicable, *p*-values and degrees of freedom are reported.

## Additional resources

Installation instructions, database management, and remote execution guidelines are available in the MMonitor README at www.github.com/lucast122/mmonitor.

## Author Contributions

T.N.L. conceived the main scientific ideas, wrote most of the software, led data analysis and interpretation, and wrote the majority of the manuscript. U.B. generated most of the experimental data and contributed to analysis and writing. A.G. contributed to scientific ideas and writing. K.G. assisted with data generation. T.L. and S.K. contributed to scientific ideas and code. R.E.L. provided scientific input. L.T.A. contributed to scientific ideas and interpretation. D.H.H. contributed to scientific ideas, interpretation, and manuscript writing, and acted as corresponding author. All authors provided feedback and helped improve the manuscript.

## Supporting information

MMonitor supplementary material

## ACKNOWLEDGMENTS

The authors thank the Angenent Lab and the Ley Lab for their sequencing data, suggestions, and tool testing. We also acknowledge members of the Algorithms in Bioinformatics labs at the University of Tübingen. Special thanks to Byoung Seung Jeon for building and sequencing the reactor that provided the WGS data. The development of MMonitor was funded through the Deutsche Forschungsgemeinschaft under Germany’s Excellence Strategy (EXC 2124 – 390838134) to REL, LTA and, DH. Additional parts of the work were funded by the Alexander von Humboldt Foundation in the framework of the Alexander von Humboldt Professorship to LTA and by the Novo Nordisk Foundation CO2 Research Center (CORC) with grant number NNF21SA0072700, published under the number CORC_25_## to LTA.

## CONFLICTS OF INTEREST

The authors declare no conflict of interest.

## Supplementary Material

### Taxonomic classification of WGS MOCK dataset

MMonitor’s pipelines use previously described methods that were already benchmarked by their authors, so we did not perform extensive benchmarks of those methods. However, since MMonitor uses updated indices for taxonomic classification, we ran it on the WGS MOCK community sample (ONT Q20 Zymo) using the same preprocessing as described in [64]. Figure **4** shows the abundances of the taxa of the MOCK community calculated by MMonitor compared to the theoretical abundances. It was generated using a python notebook from the mentioned publication [64]. MMonitor identified all genera with slight deviations in theoretical abundance only. At the species level, MMonitor correctly classified the bacterial genomes but failed to assign reads to *Cryptococcus neoformans*, which is a yeast species. Instead, it assigned the reads to a closely related fungal species of the genus *Cryptococcus*. Species-level abundances are less accurate than genus-level abundances due to more false positives summarized as Other (Fig. **4**). However, MMonitor effectively represented the sample’s taxonomic composition, aligning closely with theoretical abundance estimates.

**FIG 4.**
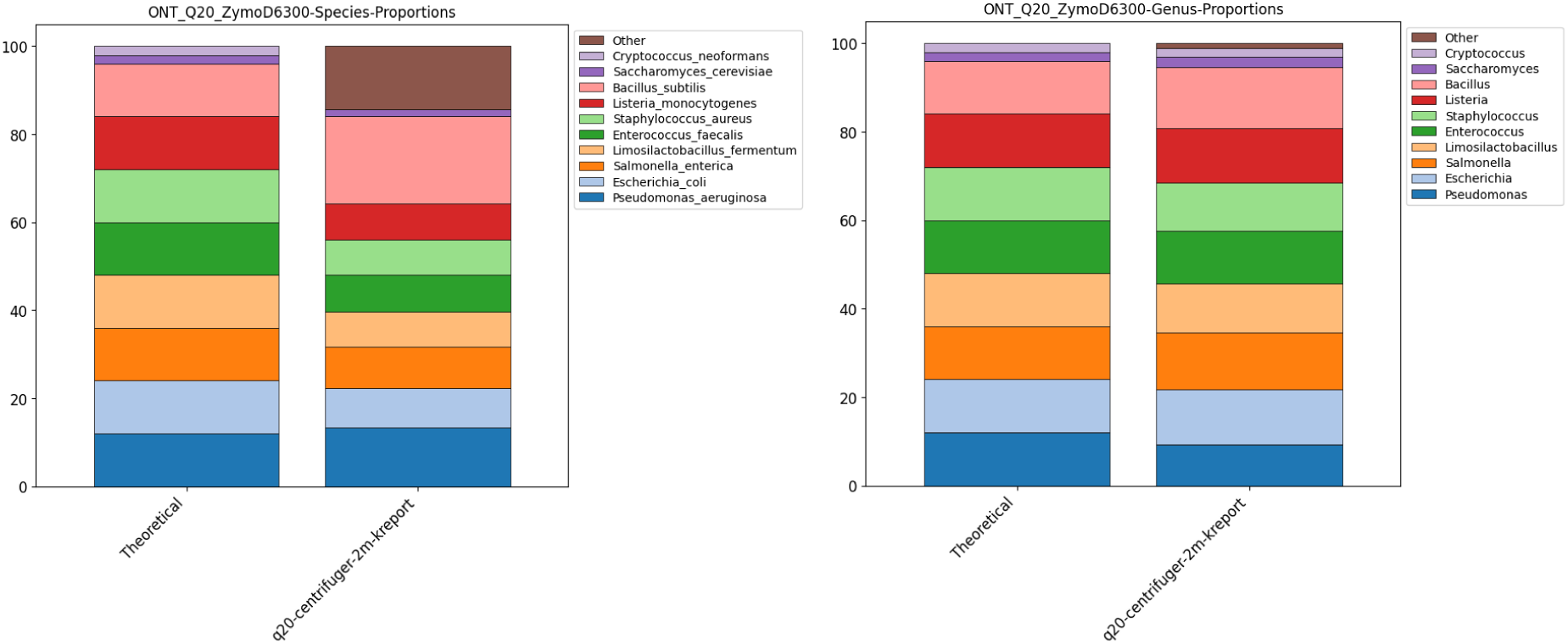
Taxonomic classification results of MMonitor on the Zymo Q20 ONT WGS mock community compared to theoretical abundances on the species level (left) and genus level (right).

### Feature comparison with less direct competitors to MMonitor

Table **3** provides a detailed comparison of MMonitor with indirect competitors that focus on broader metagenome analysis but lack real-time monitoring capabilities. Although these tools, such as MG-RAST, MEGAN, and Anvi’o, are powerful for large-scale metagenomic studies and offer deep statistical and functional analyses, they are not suited for time-sensitive workflows.

MG-RAST is an online platform for automated phylogenetic and functional analysis of metagenomic data, tailored to short read datasets which are analyzed after server upload. The service generates common plots, such as bar charts for taxonomic analysis, alongside more advanced visualizations like PCoA, rarefaction curves, and KEGG maps. In addition, it provides key diversity metrics, such as richness and evenness. However, because MG-RAST is not currently maintained and lacks a mechanism to update reference databases, there is no guarantee that all species in a metagenome will be detected accurately. We uploaded short-read data from a reactor metagenome to MG-RAST, which yielded only Genus-level identifications, restricting its utility for precise monitoring.

Although it features a sample management system, it lacks efficient multisample comparisons and time series analysis. As the service is optimized for short reads, it is unsuitable for real-time analysis and requires data uploads, leading to typical waiting times common with web-based services, a drawback for time-critical tasks like monitoring. In summary, although MG-RAST is user-friendly and offers various functionalities, its utility in monitoring metagenomes is restricted.

MEGAN [65] is a popular desktop application for taxonomic and functional analysis of metagenomic data with a GUI. Implementing numerous advanced algorithms for taxonomic and functional analysis, useful visualizations, and allowing multi-sample comparisons, it has some features useful for monitoring microbes. However, MEGAN is not optimized for real-time usage or time series visualizations. It requires previous alignment of reads against a protein reference database, which is very efficient for large amounts of input reads, but less efficient for rapid monitoring where small amounts of data are analyzed more frequently instead.

Anvi’o [66] is an open-source platform with tools for metagenomic analysis, including binning, pangenomics, and visualization, which supports customizable databases and advanced statistical methods. Anvi’o uses a command-line interface and has a steep learning curve, posing a barrier to users without computational expertise. It is not tailored for real-time monitoring or nanopore sequencing data.

QIIME2 is a powerful and extensible microbiome analysis package with a focus on interactive analysis and visualization. It provides pipelines for taxonomic classification and diversity analyses and supports a plugin architecture for extensibility. However, QIIME2 is designed primarily for amplicon (16S rRNA gene) sequencing data and is not optimized for long-read or real-time sequencing data. The command-line interface may limit accessibility for some users.

Nephele [67] is a cloud-based microbiome data analysis platform. It offers user-friendly web interfaces and integrates common microbiome analysis tools such as QIIME, mothur, bioBakery, and a5-miseq. Nephele supports both amplicon-based (e.g., 16S rRNA gene sequences) and whole metagenome shotgun sequencing data. Built on Amazon Web Services (AWS) cloud, it provides scalable computing resources, reducing the computational burden on researchers. Nephele emphasizes reproducibility by tracking all input files, configuration parameters, and virtual machine images used in data analyses.

The Pathosystems Resource Integration Center (PATRIC) [68] is a comprehensive bacterial bioinformatics resource focused on human pathogenic species. PATRIC integrates and annotates publicly available bacterial genomes using the Rapid Annotation Using Subsystems Technology (RAST) system. It provides tools for comparative genomics analysis, including genome browsing, protein family sorting, pathway analysis, and a suite of computational tools for bioinformatics analysis. PATRIC also integrates community-derived data, such as disease information, experimental data, and literature, facilitating a holistic approach to infectious disease research.

MGnify [69] (formerly known as EBI Metagenomics) is a platform provided by the European Institute of Bioinformatics (EMBL-EBI) for the assembly, analysis, and storage of microbiome sequences. MGnify offers comprehensive analysis and visualization of metagenomic data, providing access to taxonomic assignments and functional annotations for a vast number of datasets from diverse environments. The platform supports the assembly and analysis of metagenomic and metatranscriptomic datasets, including long-read sequencing technologies. MGnify employs standardized and versioned analysis pipelines, ensuring consistency and reproducibility in microbiome research.

## Notes

https://github.com/lucast122/mmonitor

### Competing Interest Statement

The authors have declared no competing interest.

