## Supplementary material for "MMonitor: Software for Real-Time Monitoring of Microbial Communities Using Long Reads": MMonitor supplementary material

### Taxonomic classification of WGS MOCK dataset

MMonitor's pipelines use previously described methods that were already benchmarked by their authors, so we did not perform extensive benchmarks of those methods. However, since MMonitor uses updated indices for taxonomic classification, we ran it on the WGS MOCK community sample (ONT Q20 Zymo) using the same preprocessing as described in [1]. Figure **1** shows the abundances of the taxa of the MOCK community calculated by MMonitor compared to the theoretical abundances. It was generated using a python notebook from the mentioned publication [1]. MMonitor identified all genera with slight deviations in theoretical abundance only. At the species level, MMonitor correctly classified the bacterial genomes but failed to assign reads to *Cryptococcus neoformans*, which is a yeast species. Instead, it assigned the reads to a closely related fungal species of the genus *Cryptococcus*. Species-level abundances are less accurate than genus-level abundances due to more false positives summarized as Other (Fig. **1**). However, MMonitor effectively represented the sample's taxonomic composition, aligning closely with theoretical abundance estimates.

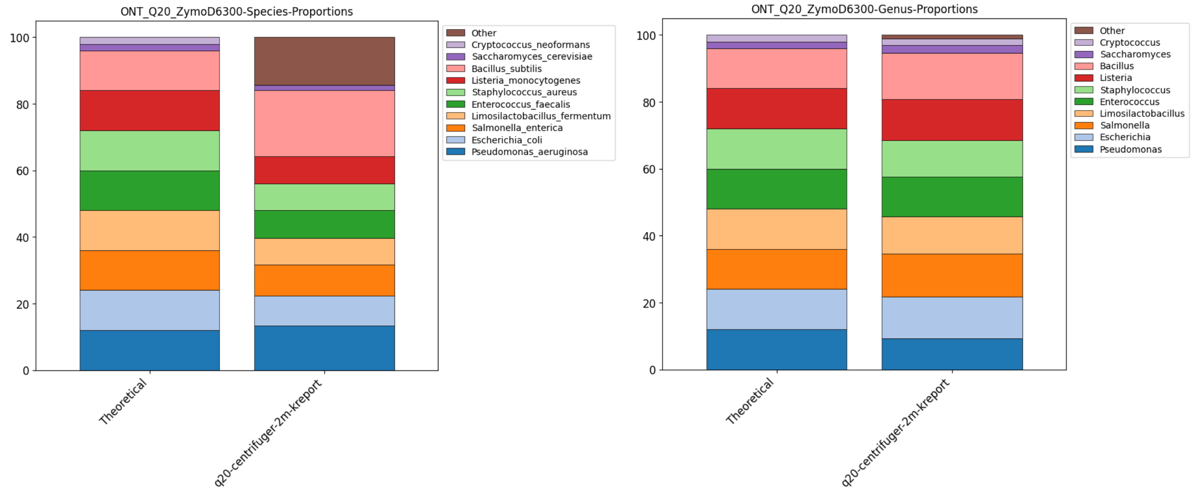


**FIG 1** Taxonomic classification results of MMonitor on the Zymo Q20 ONT WGS mock community compared to theoretical abundances on the species level (left) and genus level (right).

### Feature comparison with less direct competitors to MMonitor

**TABLE 1** Comparison of MMonitor with Indirect Competitors (Broader Metagenome Analysis Tools). N/A: Not applicable.

| Feature | MMonitor | MG-RAST | MEGAN | Anvi'o | QIIME 2 | MetaPhlAn 4 | Nephele | PATRIC | MGnify |
|---|---|---|---|---|---|---|---|---|---|
| **Time-Series Visualization** | Yes | Limited | Limited | Limited | Yes | Limited | Limited | Yes | Limited |
| **Sample Management** | Yes | Limited | No | Yes | Limited | No | Limited | Yes | Yes |
| **Speed** | Fast | Moderate | Moderate | Variable | Variable | Fast | Moderate | Moderate | Moderate |
| **Sequence Quality Control** | Yes | Yes | Yes | Yes | Yes | No | Yes | Yes | Yes |
| **Statistical Analysis Methods** | Yes | Limited | Yes | Yes | Yes | Limited | Yes | Yes | Yes |
| **User Interface** | GUI & CLI | Web-Based | GUI & CLI | GUI & CLI | CLI | CLI | Web-Based | Web-Based & CLI | Web-Based |
| **Data Accessibility** | Web, API | Web | No | No | No | No | Limited | Web | Web |
| **Customizable Databases** | Yes | No | Yes | Yes | Yes | Limited | Limited | Yes | Limited |
| **Automated Reporting** | Yes | Yes | Limited | Limited | No | Limited | Yes | Yes | Yes |
| **Scalability** | High | Moderate | Moderate | Variable | Variable | High | High | High | High |
| **Open Source** | Yes | Yes | Yes | Yes | Yes | Yes | Yes | Yes | Yes |
| **Cost** | Free | Free | Free (Academic) | Free | Free | Free | Free | Free | Free |
| **Ease of Installation** | Easy | N/A | Moderate | Moderate | Moderate | Easy | Easy | Easy to Moderate | Easy |
| **Real-Time Analysis** | Yes | No | No | No | No | No | No | No | No |
